# Low parental melatonin levels increases autism spectrum disorder risk in children

**DOI:** 10.1101/046722

**Authors:** Wiebe Braam, Friederike Ehrhart, Chris T. Evelo, Anneke P.H.M. Maas, Marcel G. Smits, Leopold Curfs

## Abstract

Low melatonin levels are a frequent finding in autism spectrum disorder (ASD) patients.
Melatonin is important for normal neurodevelopment and embryonic growth and highly effective in protecting DNA from oxidative damage. Melatonin deficiency, possibly due to low CYP1A2 activity, could be an etiologic factor. As the fetus does not produce melatonin, low maternal melatonin levels should be involved. We measured 6-sulfatoxymelatonin in urine of 60 mothers of a child with ASD that attended our sleep clinic for people with an intellectual disability (ID), and asked for coffee consumption habits, as these are known to be related to CYP1A2 activity. 6-Sulfatoxymelatonin levels were significantly lower in mothers than in controls (*p* = 0.005), as well as evening coffee consumption (*p* = 0.034).

**Abbreviations:**

ASD: Autism spectrum disorder
ID: intellectual disability
6-SM: 6–sulfatoxymelatonin

## Introduction

Autism Spectrum Disorder (ASD) refers to a heterogeneous group of neurodevelopmental disorders, with a complex multifactorial etiology, starting very early in life. The disease is characterized by abnormal social interactions, impaired language development, and stereotyped and repetitive behaviours. ASDs have in common that they each share one or more core symptoms of autism. But at the same time there are large differences between them, both with regard to symptomatology as to etiology (Parellada et al. 2014). Intellectual disabilities (ID) and ASD, although apparently two separate disorders, co-occur much more frequent in the same patient than by chance. While in 40% of persons with ID autism is present, at the same time 70% of people with autism also has an intellectual disability (Matson and Shoemaker 2009). The frequent co-occurrence of ASD and ID, however, raises questions whether this is a true co-occurrence because both share, at least in part, similarities in their etiology. Especially since in more severe cases of ID, the risk of ASD is greater (Vig and Jedrysek 1999), as is the risk of ID in more severe cases of ASD (Gotham et al. 2012).

There is a growing number of literature on the genetic basis of ASD. Over a hundred of chromosomal abnormalities have been found, however, not one of them is responsible for more than 1% of ASD cases. This means that the underlying genetic process in ASD is complex. Recent reviews (Kim and Leventhal 2015; Li et al. 2012; X. Liu and Takumi 2014; Rosti et al. 2014; Sanders et al. 2015) provide a good overview of the current state of knowledge.

The relative risk of a newborn child from parents having a child with ASD is 25-fold increased compared to the general population, suggesting a genetic component (Li et al. 2012). However, in only about 10% a specific genetic disorder is diagnosed (X. Liu and Takumi 2014). It is assumed that ASD is caused by a combination of *de novo* inherited variation and common variation. In a recent epidemiological study in Sweden, it was possible to distinguish between both components, estimating liability of *de novo* variations to be 2.6% and common variation to be 49% (Gaugler et al. 2014). In a recent large study in twins with ASD, concordance for monozygotic male twins was 77% (females 50%), and for dizygotic male twins 31% (females 36%) (Hallmayer et al. 2011). Based on their data they were able to calculate that the contribution of genetic factors was smaller (38%) than shared environmental factors (58%).

Epidemiological studies have identified several environmental factors that increase ASD risk (Gardener et al. 2009, 2011; Polo-Kantola et al. 2014; Kuzniewicz et al. 2014; Walker et al. 2015) (see supplementary file). Most important factors are advanced parental age at birth, bleeding, gestational diabetes, having a mother born abroad, fetal distress, birth injury or trauma, multiple birth, maternal haemorrhage, summer birth, low birth weight, small for gestational age, congenital malformation, low 5-minute Apgar score, feeding difficulties, meconium aspiration, neonatal anaemia, ABO or Rh incompatibility, and hyperbilirubinemia. Also subnormal breast feeding increases autism risk (Al-Farsi et al. 2012). Interestingly, many of these environmental factors can be linked to melatonin deficiency.

Disturbances in melatonin turnover leading to different melatonin levels in ASD are found in several studies (Rossignol and Frye 2011; Tordjman et al. 2013; Veatch et al. 2015). Melatonin levels in ASD are negatively correlated with severity of autistic impairments (Tordjman et al. 2005).

Melatonin is a hormone with multiple functions and it is synthesized in the pineal gland during the dark phase of the day and involved in sleep induction and circadian rhythm regulation (Arendt 2005). Melatonin also acts as a free radical scavenger and antioxidant and is highly effective in reducing effects of oxidative stress throughout the body (Bejarano et al. 2014; Reiter et al. 2014; Korkmaz et al. 2012). There are also indications that melatonin protects against oxidative stress-evoked DNA damage (Bejarano et al. 2014; Ferreira et al. 2013; Reiter et al. 2014; Reiter et al. 2007).

Melatonin is synthesized from serotonin in a two step process, involving the enzymes AANAT and ASMT (see Figure 1) (Ackermann and Stehle 2006). The expression of their mRNAs encoding for these enzymes follows a day/night rhythm.

**Figure 1.**
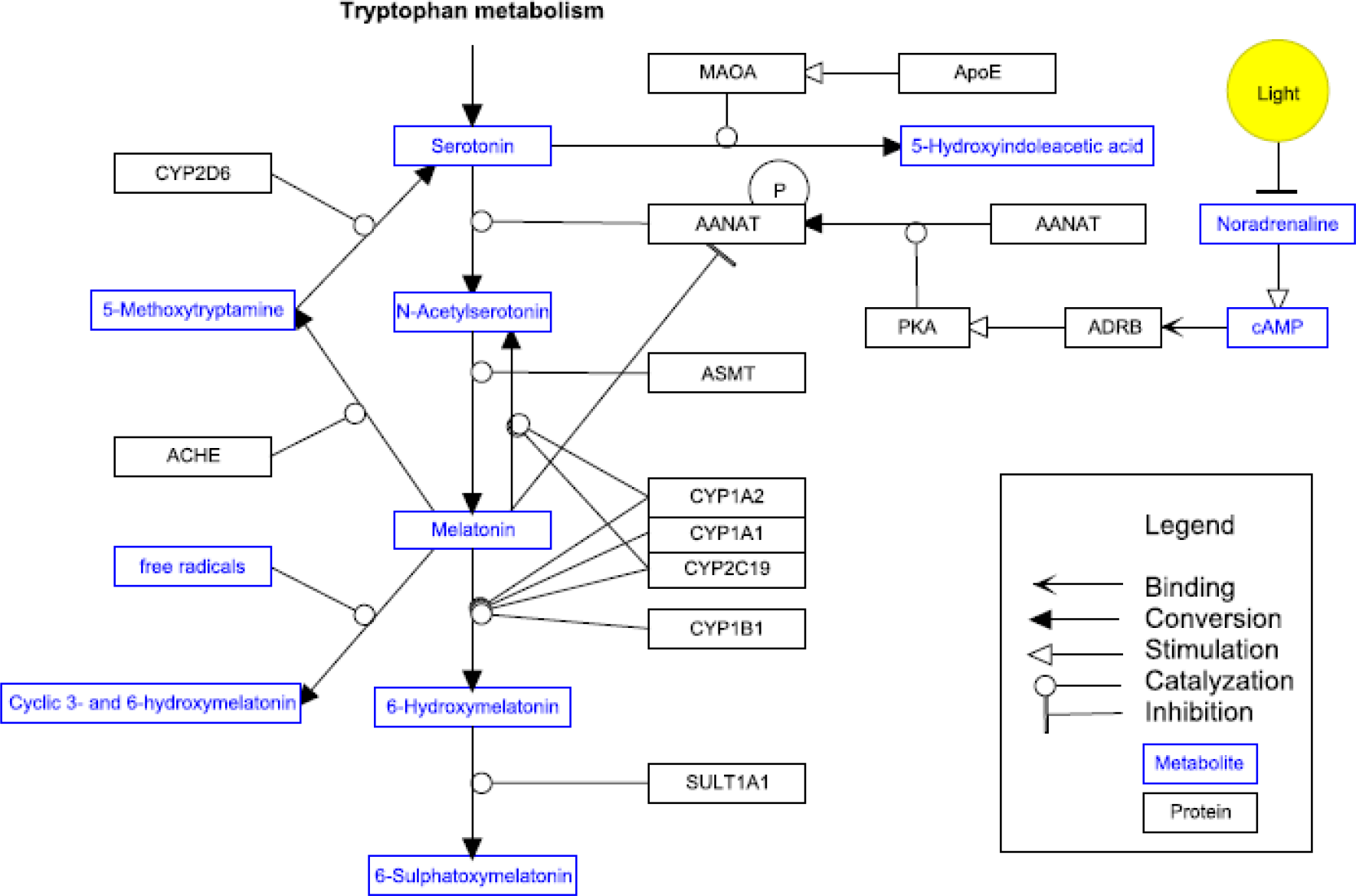
Synthesis and breakdown pathway of melatonin. Figure modified from http://www.wikipathways.org/instance/WP3298

AANAT is generally considered to be the rate-limiting enzyme in the melatonin synthesis pathway. However, recent studies indicate that the rate limiting role at night is partly taken over by ASMT (T. Liu and Borjigin 2005). Exposure to light rapidly decreases AANAT activity. Timing, profile and amount of melatonin secretion display large inter-individual differences, but are in healthy individuals highly reproducible from day to day like a hormonal fingerprint (Arendt 2005). As information on normative data of melatonin levels is scarce and internationally accepted normative data are not available, it is also not clear which melatonin levels are pathologically low. This is remarkable, given the large number of functions of melatonin. Although several positive functions are attributed to melatonin, melatonin deficiency is not regarded as a risk factor or a disease yet.

Melatonin is metabolized principally by 6-hydroxylation to 6-hydroxymelatonin, which is sulphated to 6-sulfatoxymelatonin (6-SM) and excreted in urine (Ma et al. 2005; Yu et al. 2003) (Figure 1). CYP1A2, and to a lesser extend CYP1A1 and CYP2C19 are the main cytochrome isozymes to be responsible for the plasma clearance of melatonin in the liver (Ma et al. 2005). CYP1A2 activity in fetal and neonatal liver tissue is very low to undetectable (Sonnier and Cresteil 1998). The expression of CYP1A2, as well as other members of the cytochrome P450 gene family, is regulated i.a. by the biological clock (Froy 2009). Melatonin does not influence CYP1A2 activity at physiologic levels. Only at extremely high concentrations of 3–300 µM an inhibition was observed (Chang et al. 2010).

The half life of exogenous melatonin ranges between 28 to 126 minutes (Fourtillan et al. 2000; Harpsoe et al. 2015). Differences in melatonin half-life reflect the wide inter-individual differences (10 to 200-fold) in CYP1A2 activity (Gunes and Dahl 2008). Several reports indicate that SNPs in the CYP1A2 gene are associated with increased inducibility, decreased activity or inducibility or even loss of activity of the CYP1A2 enzyme as compared to the wild type CYP1A2*1A (Sachse et al. 1999; Zhou et al. 2009a; Zhou et al. 2009b; Zhou et al. 2010; Nakajima et al. 1999; Chevalier et al. 2001). The (sub)variant alleles associated with decreased of absent activity of CYP1A2 are *1C, *1K, *3, *4, *5, *6 and *7. The proportion of individuals with the slow phenotype varies among ethnic populations (Zhou et al. 2010).

Caffeine clearance is considered as the gold standard for assessment of CYP1A2 activity, because more than 90% of the primary metabolism of caffeine depends on CYP1A2 (Hartter et al. 2006). Caffeine related side effects, i.e. caffeine-induced insomnia, are more frequent in people who are CYP1A2 slow metabolizer (Carrillo and Benitez 1996). Several polymorphisms decrease the metabolic activity of CYP1A2 and are significantly inversely related to coffee intake, so coffee intake is influenced by CYP1A2 genotype (Josse et al. 2012; Rodenburg et al. 2012). In people with caffeine-induced insomnia caffeine clearance is significant slower than in people without sleep onset problems after drinking coffee in the evening (Levy and Zylber-Katz 1983).

### Melatonin and its relation to ASD

Low levels of melatonin are found in the majority of individuals with ASD (Rossignol and Frye 2011; Tordjman et al. 2005; Nir et al. 1995; Kulman et al. 2000). Studies on genes involved in melatonin metabolism have long been focussed on the melatonin synthesis pathway, mainly of ASMT. Interestingly no studies were performed on the melatonin catabolism by CYP1A2 until in 2013 a possible relationship between slow CYP1A2 polymorphisms in slow melatonin metabolizers and ASD was proposed (Braam et al. 2013). Even though ASD frequently is associated with low melatonin levels and suspicion exists for an involvement of the ASMT gene in the etiology of autism, several studies could not find any evidence for a significantly higher incidence of abnormalities of the ASMT gene in ASD (Melke et al. 2008; Pagan et al. 2014). And if so, ASMT abnormalities were also found in healthy controls.

Another potential link between melatonin and ASD etiology was proposed by Braam et al. in an earlier report (Braam et al. 2013). In a study in 15 consecutive patients with ID and sleep problems, presenting with disappearing effect of melatonin treatment after an initial good response, extremely high (>50 pg/ml) melatonin levels at noon and 04:00 pm were found. They hypothesized that the disappearing effectiveness of melatonin treatment was associated with slow metabolization of melatonin due to a SNP of CYP1A2 and analysed DNA for a CYP1A2 SNP. In 8 patients a SNP was found. Because 13 of 15 patients with CYP1A2 slow metabolism were diagnosed with ASD, it was hypothesized that low melatonin levels in autism are caused by a yet unknown mechanism in which CYP1A2 slow metabolism is involved, and that low melatonin levels are possibly part of the mechanisms that <u>*cause*</u> ASD, rather than that low melatonin levels are the <u>*result*</u> of ASD.

This potential mechanism was investigated by Veatch et al. (2014) (Veatch et al. 2014). They found a higher frequency of dysfunctional ASMT polymorphisms than earlier reported (Melke et al. 2008; Toma et al. 2007; Jonsson et al. 2010; Wang et al. 2013). They also found substantially higher rates of CYP1A2 polymorphism in ASD patients than in the previous report (Braam et al. 2013). Based on the linkage disequilibrium structure across SNPs in ASMT and CYP1A2 in their subset of individuals, Veatch et al. predicted a potential gene-gene interaction (Veatch et al. 2015).

The embryo and fetus do not secrete melatonin and are totally dependent on maternal melatonin (Merchant et al. 2013; Kennaway et al. 1996). Melatonin secretion begins no earlier than approximately 9 - 15 weeks after birth, depending of gestational age (Kennaway et al.1996). Maternal melatonin passes the placental barrier easily and the fetal circadian melatonin rhythm follows the maternal rhythm (Reiter et al. 2013). Maternal melatonin is also passing easily into breast milk, and levels are a reflection of the circadian melatonin rhythm of the mother (Cohen Engler et al. 2012). ASMT and AANAT gene activity and melatonin levels in disturbed pregnancies (intra uterine growth retardation and in severe preeclamptic women) are lower than in normal pregnancies (Tamura et al. 2008; Lanoix et al. 2012).

Melatonin is important for normal neurodevelopment and embryonic growth and helps regulate synaptic plasticity (Kong et al. 2008; Merchant et al. 2013; Rossignol and Frye 2011; Tordjman et al. 2013). Melatonin is implicated in neural growth and synapse formation and it can modulate neurite outgrowth in cultured neuronal cells (Jonsson et al. 2010). Melatonin guides the differentiation of neuronal stem cells (*in vitro)* as follows (Kong et al. 2008): It stimulates the mRNA and protein levels of TH, which is a marker for dopaminergic neurons. Dopaminergic neurons are neurons which are able to release dopamine and have therefore depending on their location in the brain 1) a high impact on regulation of cognitive function, emotional behaviour, and reward system or 2) regulate motor function. *In vivo,* melatonin increases dendritogenesis (dendrite formation, enlargement and complexity) in the hilar and mossy neurons of hippocampus in rats (Dominguez-Alonso et al. 2015). The accompanying *in vitro* results indicate that melatonin triggers CALM which stimulates MAP2 via CAMK pathway (Figure 2). Melatonin increases the total cellular level of CALM and fosters the translocation from cytosol to membrane/cytoskeleton interfaces (Dominguez-Alonso et al. 2015).

**Figure 2.**
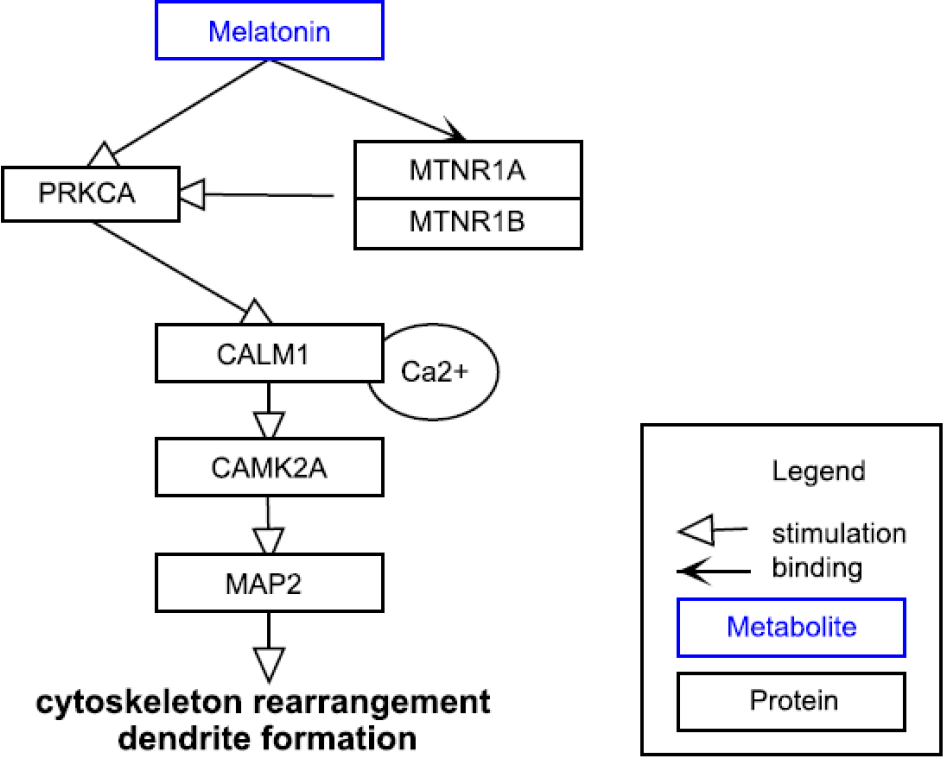
Melatonin influences the cytoskeleton arrangement and dendrite formation in neuronal cells via CALM and CAMK. Figure modified from http://www.wikipathways.org/instance/WP3298

Although the genetic basis of ASD is not disputed, no single chromosomal abnormality is responsible for >1% of ASD cases. Low melatonin levels are found in the majority of ASD cases, but is regarded as co-morbidity and not considered as part of ASD etiology. All data mentioned above can be linked directly or indirectly with melatonin. As ASD is already present at birth and the fetus does not produce melatonin, low maternal melatonin levels should also be involved. Low melatonin levels are not only caused by disturbances in melatonin synthesis, but are likely also related to low CYP1A2 activity, in view of the finding of CYP1A2 gene SNPs in ASD patients and a potential gene-gene interaction between CYP1A2 and ASMT. We therefore measured 6-SM in urine of mothers of a child with ASD that attended our sleep clinic, and asked for parental coffee consumption habits, as these are known to be related to CYP1A2 activity.

## Methods and Materials

### Participants

Approval was obtained through the Ethical Committee of [blinded for review]. Mothers of patients with chronic sleep disturbance, who visited our sleep centre for people with ID between January and July 2015, were asked to participate if her child was diagnosed with ASD. Mothers of children with a genetic ID syndrome and an autistic phenotype were also asked to participate. Out of 97 children seen in this period, 18 were not diagnosed with ASD and 13 mothers were not eligible for several other reasons (five children were adopted or foster child, four children were only seen once, with three mothers we had severe language barriers, and one mother used melatonin supplementary on a daily basis). So, 66 mothers consented to participate. The control group consisted of 15 women who work in the building where the sleep centre is located. These women all had healthy children without signs of autism and siblings of controls also did not have signs of autism. All participants were of Caucasian origin.

### Procedure

After explaining purpose and procedure of the study, participants received an introductory letter with information of the study and an informed consent letter to be send back by return paid mail. After receiving the signed consent letter, a questionnaire, together with a shipping container for a urine sample and instruction sheet was send to participants. Participants were instructed to empty the bladder at bedtime. Time of voiding at bedtime and first voiding in the morning were recorded. The first morning urination, plus any urine voided during the night, was collected and the total urine volume was recorded. A sample of the total urine was poured into a plastic bottle and sent to the sleep centre where it was frozen and stored at minus 20°C until measurements took place. The questionnaire (see supplementary data) contained a total of 17 questions, namely six questions on age, weight, medication use, and number and age of children (oldest and youngest child), seven questions based on the Insomnia Severity Index (Bastien et al. 2001) and four questions about the use of coffee.

### 6-SM concentration measurement

6-SM in urine was analysed by isotope dilution mass spectrometry using online solid phase extraction in combination with liquid chromatography and tandem mass spectrometry (XLC-MS/MS) (Van Faassen et al., manuscript in preparation). In short, 100 µl of urine was used for the analysis, deuterated 6-SM was used as internal standard. Mean intra-and inter-assay coefficients of variation were below 6.0%. Quantification limit for 6-SM was 0.2 nmol/l.

As advanced parental age at birth is a factor associated with autism risk (Gardener et al. 2009) and melatonin levels decrease with 2.7% per year (Kennaway et al. 1999), we recalculated the urinary 6-SM results in mothers with an ASD child, taking into account the age at which they received this child. In 6-SM levels in controls, we based recalculating on their age at receiving their first-born child. In case controls had more than one child, we took the mean of their age receiving their oldest and their youngest child.

### Statistical analysis

All statistics were performed using IBM SPSS software (version 20, SPSS Inc., Chicago, IL, USA). T-tests for independent samples were conducted to test differences between ASD-groups and controls. To test for differences on sleep questions between the subgroups non-parametric tests (Fisher's exact test and Mann-Whitney test) were used because of low cell frequencies and/or not normally distributed data. To test the relationship between sleep and melatonin bivariate correlation (Kendall's tau-b) was used. Statistical significance was accepted at *p* < 0.05.

## Results

Of 66 mothers, who consented to participate in this study, one withdrew without specifying a reason. In five cases the urine samples did not arrive. So urine samples of 60 mothers were analysed. Their mean age was 42.9 (± 5.7) years. The mean age of 15 women in the control group was 44.3 (± 9.7) years. No significant differences with respect to age (*p* = 0.486), weight (*p* = 0.270) and time in bed (*p* = 0.985) were found between participant and control group. However, mean age of mothers of an ASD child at time of delivery was higher (31.2 years) than the mean age control mothers received their children (28.1 years). This difference was significant (*p* = 0.01). Other demographic data are shown in Table 1:.

**Table 1.** Demographic data

| Demographic data | ASD | | Control | | $p$ |
| --- | --- | --- | --- | --- | --- |
|  | Mean | SD | Mean | SD |  |
| Age of mother at study participation | 42.92 | 5.72 | 44.27 | 9.73 | 0.486 |
| Age of mother at birth child | 31.17 | 4.15 | 28.07 | 3.58 | 0.01 |
| Number of children | 2.18 | 0.89 | 2.40 | 0.51 | 0.37 |
| Weight of mother | 73.24 | 13.04 | 69.27 | 8.23 | 0.27 |
| Time in bed | 8.01 | 0.12 | 8.01 | 0.18 | 0.98 |

Data on etiological diagnoses of children with ASD are presented in (Table 2. It is noteworthy that there are substantial groups with Angelman syndrome or Smith Magenis syndrome in the group studied. It should however be noted that both syndromes occur relatively frequently (prevalence 1:20.000) and that they are well known for their sleep problems (Dagli et al. 2012; De Leersnyder 2006).

**Table 2.** Etiological diagnoses of ASD children

|  |  |  |
| --- | --- | --- |
| Angelman syndrome | 10 | (group AS) |
| Smith Magenis syndrome | 18 | (group SMS) |
| Other diagnoses | 22 | (group Other) |
| 6p25 deletion | 2 |  |
| 15q13.2 duplication | 1 |  |
| 16p11.2 deletion | 2 |  |
| 16p11.2 duplication | 1 |  |
| 16p13.3 deletion | 1 |  |
| 17p11.2 duplication (Potocki syndrome) | 1 |  |
| 17q21 deletion and 16q22.1 deletion | 1 |  |
| 18q12.1 deletion | 2 |  |
| 22q11 duplication | 1 |  |
| ADNP gene mutation | 1 |  |
| Congenital cytomegaly | 1 |  |
| Down syndrome | 1 |  |
| Isodicentric chromosome 15 syndrome | 1 |  |
| Phelan McDermid syndrome | 2 |  |
| SCN2A gene mutation | 1 |  |
| SOX21 and GPC6 gene mutation | 1 |  |
| Translocation 10:22 (der(10)t(10:23)(qter;qter) | 1 |  |
| Williams syndrome | 1 |  |
| Unknown | 10 | (group Unkown) |
| Total | 60 |  |

In this study we found a significant lower urinary 6-SM excretion in mothers of a child with ASD compared to control mothers (*p* = 0.012). Group means and SDs of the excretion of 6-SM excretion in the urine in all groups are given in Table 3. Melatonin levels are known to decline after the age of 20 years with 2.7% (Kennaway et al. 1999) each year. Because ages of mothers and children were asked for, we were able to recalculate 6-SM excretion levels in each mother to the year their child was born. Comparing urinary 6-SM levels in the year their child was born is more relevant, given our hypothesis that low melatonin levels in pregnancy are a risk factor for getting a child with ASD. As shown in Table 3 differences in recalculated 6-SM excretion levels between mothers of a child with ASD compared to control mothers were significant (*p* = 0.002).

**Table 3.** 6-SM/creatinine ratio in urine measured and recalculated to the age of the mother at child birth

|  | measured |  | recalculated |  |
| --- | --- | --- | --- | --- |
|  | ASD | Control | ASD | Control |
| **n** | 60 | 15 | 60 | 15 |
| **6-SM/creatinine** | 7.67±3.62 | 10.42±6.72 | 10.52±5.20 | 16.67±10.72 |
| **Min - Max** | 1.99 – 16.25 | 2.31 – 25.5 | 2.81 - 21.93 | 4.79 - 42.59 |
| **p** | 0.012 |  | 0.002 |  |

**Figure 3.**
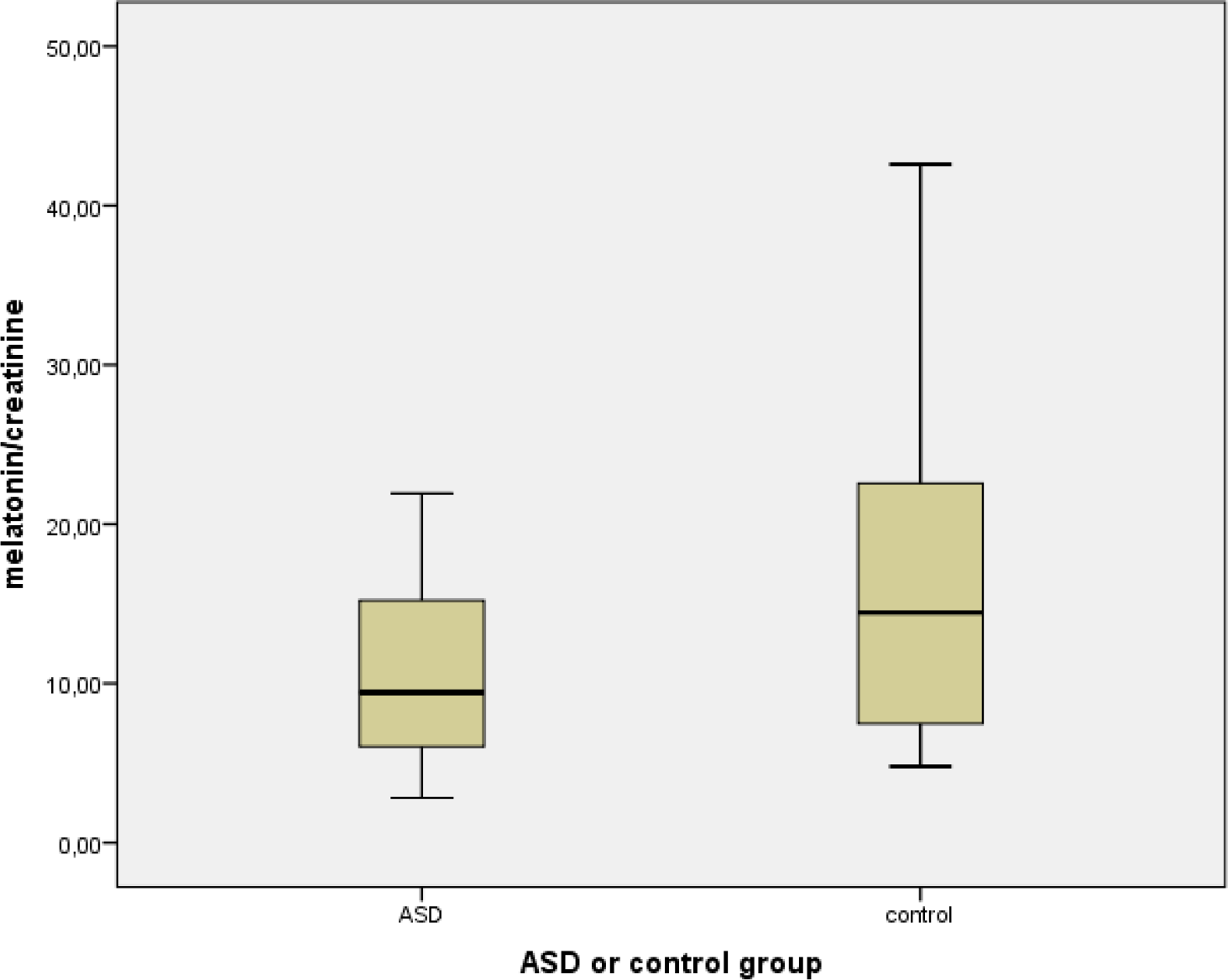
6-SM/creatinine ratio in urine of mothers. The values have been recalculated to the age of the
mother at child birth.

Looking into subgroups, only mothers in the group with other genetic disorders and with Angelman syndrome had a significant lower urinary 6-SM excretion compared to control mothers (*p* = 0.012 resp. 0.042) (Table 4). Urinary 6-SM levels in mothers of a child with Smith Magenis syndrome also were lower than in controls, and these differences approached significance (*p* = 0.05).

**Table 4.** 6-SM/creatinine (nmol/l) in urine (recalculated to age of mother at child birth) for different ID groups; ASD eci (no genetic disorder found yet), ASD syndromal (ASD in child with a chromosomal disorder, but not Angelman of Smith Magenis syndrome), Angelman syndrome and Smith Magenis syndrome.

|  | ASD eci | ASD syndromal | Angelman syndrome | Smith Magenis syndrome | Controls |
| --- | --- | --- | --- | --- | --- |
| **n** | 10 | 22 | 10 | 18 | 15 |
| **6-SM/creatinine** | 13.37±5.20 | 9.54±5.63 | 8.92±4.46 | 11.03±4.70 | 16.67±10.72 |
| **Min - Max** | 4.96 - 21.55 | 3.33 - 21.93 | 4.46 - 19.91 | 2.81 - 18.77 | 4.79 - 42.59 |
| **p** | 0.375 | 0.012 | 0.042 | 0.052 |  |

Five mothers of a child with ASD also had another child with ASD and/or an ID. These children, however, were not treated for sleep problems in our sleep centre. One child was known with Prader Willi syndrome, being genetically of paternal origin. Melatonin levels in the other four mothers were lower (mean 6.54 nmol/l ± 3.04) compared to the other mothers with one child with ASD (mean 10.80 nmol/l ± 5.22, *p* = 0.058), but this was not significant due to low numbers.

The number of mothers drinking coffee does not differ between mothers with an ASD child and controls (*p* = 0.375). However, more mothers with an ASD child do not drink coffee after 06:00 pm than mothers in the control group (*p* = 0.018). Time of the last cup of coffee was significantly later in control mothers than in mothers with an ASD child (07:10 pm, resp. 05:05 pm: *p* = 0.003) (Figure 4). Urinary 6-SM levels in mothers that do not drink coffee after 06:00 pm were significantly lower (13.79 nmol/l ± 9.08) than in mothers that do drink coffee after 06:00 pm (10.39 nmol/l ± 4.90) (*p* = 0.039). Also more fathers with an ASD child do not drink coffee after 06:00 pm (*p* = 0.020).

**Figure 4.**
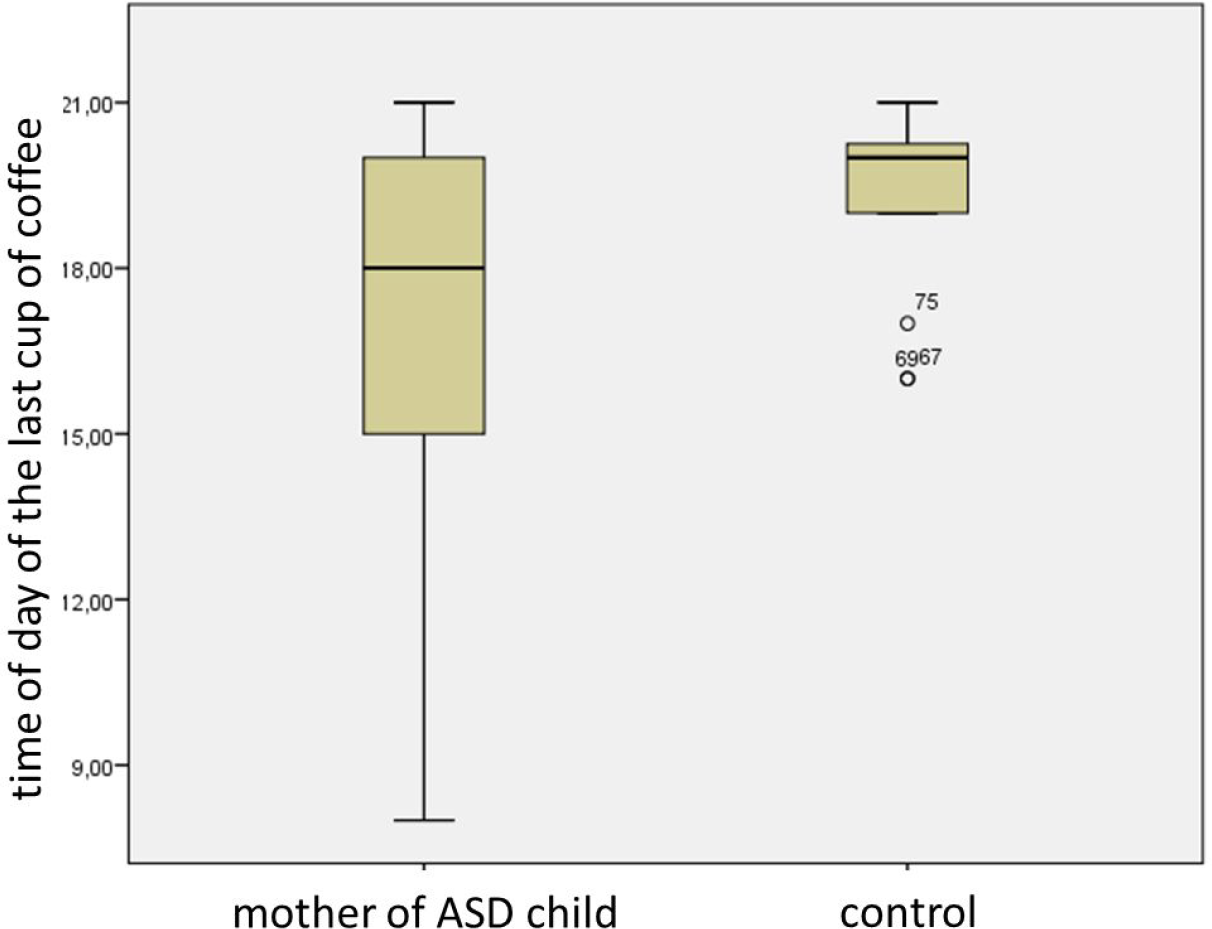
Time of the day of the last cup of coffee for mothers of an ASD child and control.

Of 60 mothers who‘s urine samples were analysed, one did not fill in the questions on sleep. So questions on sleep of 59 mothers of the patient group and 15 mothers of the control group were analysed. If a mother indicated that she did not experience difficulty with either falling asleep or staying asleep and did not have a problem with waking up too early she was asked to skip three questions and no total score on the Insomnia Severity Index (ISI) could be calculated. A cut-off score of 10 was used to indicate clinical significant insomnia (Morin et al. 2011). Of the women with an ASD child (n = 33) 67% met the cut-off score of the ISI compared to 29% of the women in the control group (n = 7). This difference was not statistically significant (*p* = 0.094, Fisher’s exact test).

In order to compare sleep between all mothers of the patient and the control group we also calculated a shortened ISI (sISI) based on two questions (severity of sleep problems and satisfaction with current sleep pattern) filled in by all mothers (n = 74). Higher scores indicate more severe sleep problems and less satisfaction with current sleep pattern. Scores on sISI in patient group (Mdn = 4) did not differ significantly from the control group (Mdn = 2), U = 542.50, z = 1.366, p = 0.175, r = 0.16. Furthermore no correlation was found (tau = 0.00) between sISI and the excretion of 6-SM in the urine (*p* = 0.996, two-tailed).

## Discussion

In this study we found a significant lower 6-SM excretion in mothers of a child with ASD. This difference cannot be explained by the difference in their age at birth of their child.

ASD is caused by a mixture of rare and common genetic variation, as well as environmental factors. Melatonin deficiency can play a role in all three of these factors. Epidemiological studies have identified several environmental factors that increase ASD risk. Almost all of the known prenatal, perinatal and neonatal ASD risk factors, found in two metaanalyses (Gardener et al. (2009) (Gardener et al. 2009) and Gardener et al. (2011) (Gardener et al. 2011)) can directly or indirectly be linked to melatonin deficiency (see supplementary files). Additionally, low CYP metabolization indicated by coffee drinking behaviour was found to be an early indicator. A correlation between ASD and low melatonin levels/slow CYP metabolizers has been found earlier in different experimental settings (see (Table 5).

**Table 5.** Experimental and population data which shows correlation of ASD and low melatonin/CYP1A2

|  | **Correlation of ASD and melatonin/CYP data** |
| --- | --- |
| **Population** | Prevalence estimates of ASD ranges from 30.0 : 10 000 to 116.1 : 10 000. Lower prevalence estimates are found in Australia (39.2 : 10 000) and China (11.6 : 10 000), and the highest rates came one Japanese study (181.1 : 10 000) (Elsabbagh et al. 2012).<br>Slow metabolizer phenotype, using substrate probes, has been identified in Australians (5%), Japanese (14%) and Chinese (5%), while Asian and African populations have lower CYP1A2 activity compared with Caucasians (Zhou et al. 2009b). See also sup. data Table 1. |
| **Gender difference** | Advanced genetic research did not yet identify chromosomal loci that contribute to the gender difference. Perhaps the gender difference in melatonin levels can offer an explanation: Melatonin levels in girls are two to threefold higher than in boys (Fideleff et al. 2006). A theoretical explanation could be that the damage to fetal brain development by low maternal pregnancy melatonin levels, after birth may be limited in case melatonin production in the child is normal. As melatonin levels in girls are two to threefold higher than in boys (Fideleff et al. 2006), this could ultimately lead to less, or much milder cases of autism in girls than in boys. After puberty the gender difference is smaller. Melatonin amplitude in women is one and a half times higher than in men (Cain et al. 2010). |
| **Mouse model** | Investigation of melatonin supplementation in the BALB/cJ inbred strain of mice on behaviour in off-spring, as well on brain morphology, is advised as this mice strain is already an ASD mouse model because it displays all core behavioural features of ASD (Blanchard et al. 2013; K. Z. Meyza et al. 2013; K. Meyza et al. 2015). This mice strain happens to be melatonin deficient (Vollrath et al. 1988). |

Finding a significant lower urinary 6-SM excretion in mothers of a child with ASD compared to control mothers, combined with the neurotrophic properties of melatonin, total dependence of the fetus on maternal melatonin and a link between already established ASD risk factors and melatonin, supports our hypothesis that low melatonin levels may be the shared “environmental factor” postulated by earlier twin studies (Hallmayer et al. 2011).

Low endogenous melatonin levels in slow melatonin metabolizers seems to be a contradiction, because there are several publications in which not low, but high melatonin levels were found in connection to CYP1A2 slow metabolism. However, it should be clear that in these studies artificial situations were created, in which melatonin levels were measured. First, when exogenous melatonin is taken in combination with caffeine, serum melatonin levels are higher than without caffeine coadministration (Hartter et al. 2003). This situation, however, does not describe endogenous melatonin levels, but the effect of a combined caffeine and exogenous melatonin intake on serum melatonin levels. Second, ingestion of fluvoxamine increases serum melatonin level by inhibiting CYP1A2 activity in healthy subjects (von Bahr et al. 2000). This situation, however, also does not describe endogenous melatonin levels, but the effect of inhibiting CYP1A2 activity by fluvoxamine in subjects with otherwise normal CYP1A2 activity. And third, in subjects who are CYP1A2 slow metabolizer, salivary melatonin levels are higher after exogenous melatonin intake than in subjects who are not a CYP1A2 slow metaboliser (Braam et al. 2010; Braam et al. 2013). Also this situation does not describe endogenous melatonin levels, but the effect of exogenous melatonin in subjects in which CYP1A2 activity is low.

Strong evidence points at a substantial contribution of common genetic variation to ASD (Gaugler et al. 2014). Low CYP1A2 activity is a common heritable genetic variation which can cause low ASMT transcript production (Veatch et al. 2015) and by that low melatonin levels. Additionally, the SNP reduced enzyme kinetics of CYP1A2 may show substrate saturation or inhibition effects at much lower concentration as wild type. For CYP1A2 and melatonin this has not been measured before and remains speculative. The dual effect of melatonin (neurotropic properties and protection against oxidative DNA damage), provides an explanation for multiple questions about the complex underlying genetic process in ASD. This may explain results of a recent study in quartet families with ASD (parents and two ASD-affected sibs) whole-genome sequencing, which revealed that many (68.4%) of the affected siblings did not share the same mutations, but carried different ASD-relevant mutations (Yuen et al. 2015). It also may explain why in 40% of persons with intellectual disabilities autism is present, and at the same time 70% of people with autism also has an intellectual disability (Matson and Shoemaker 2009). Our finding that in mothers of a child with ASD, who also have another child with ASD and/or an ID, urinary 6-SM excretion was lower compared to mothers with one child with ASD, is in line with these findings. Low parental melatonin levels can act on two seemingly independent levels: 1) directly by causing disturbances in brain development of the child and 2) indirectly by increasing risk of DNA damage in maternal and paternal germ cells. If melatonin levels in the child with autism are also low, this contributes to greater influence during the further development of the brain.

In a large study, aimed to find normative data on the melatonin excretion in urine, overnight urinary melatonin excretion values in healthy individuals ranged from 0.023 to 0.842 nmol/l and correlated inversely with age (Wetterberg et al. 1999). Melatonin levels did not have a Gaussian distribution, but showed a bimodal distribution, distinguishing two groups of individuals: low and high melatonin excretors.

There is a decrease in time with 25% in healthy adults aged 36 to 50 years, compared to those in the age group 20 to 35 years (Mahlberg et al. 2006; Kennaway et al. 1999). Kennaway et al. (Kennaway et al. 1999) calculated the yearly decline in melatonin plasma peak levels at night between young age groups (21 to 33 years) and old age groups (49 – 85 years) to be 2.7%, based on data in 11 studies. This is the reason why we decided to offer the measured and the recalculated data (Table 3) taking the postulated age dependant melatonin decrease into account.

Caffeine and melatonin are primarily metabolized by CYP1A2. We found that significantly more mothers with an ASD child than in the control group do not drink coffee after 6:00 pm (*p* = 0.034), mostly in order to prevent coffee induced in (*p* = 0.039). This is in line with the suggested mechanism that reduced CYP1A2 metabolic activity is connected with lower levels of ASMT transcript production (Veatch et al. 2014) and supports our hypothesis that CYP1A2 slow metabolism due to a CYP1A2 SNP is involved in low melatonin levels in autism ethiology (Braam et al. 2013). As earlier studies already have shown that caffeine clearance by CYP1A2 is significant slower in people with caffeine-induced insomnia (Levy and Zylber-Katz 1983), and that metabolic activity of CYP1A2 is significantly inversely related to coffee intake (Josse et al. 2012; Rodenburg et al. 2012), we hypothesize that coffee drinking habits can serve as an indicator for low CYP1A2 activity as well as for low melatonin levels.

Our finding that significantly more fathers with an ASD child than in the control group do not drink coffee after 6:00 pm suggests that they also could be a CYP1A2 slow metabolizer and by that have low melatonin levels. This finding is important and can explain why 76% of the *de novo* mutations, found in a study in quartet families with ASD (parents and two ASD-affected sibs), were of paternal origin (Yuen et al. 2015).

The limitation of our study is that we did not measure CYP1A2 activity, but relied on information on parental coffee consumption habits, as these are known to be related to CYP1A2 activity, and that our control group is not sufficiently large to draw firmer conclusions. Our pilot study needs to be replicated on a much larger scale. It should involve melatonin levels and CYP1A2 activity measurements in ASD patients, as well as both of their parents. The outcomes of this study implicate that in yet to be determined risk groups, melatonin supplementation may be a logical consequence. Nevertheless, melatonin supplementation in pregnant and new-borns that are at risk may possibly enhance brain development but needs careful investigation of possible risks.

This is to our knowledge the first experimental evidence of low melatonin levels in mothers being a risk factor for child ASD. It is a comprehensive theory on ASD etiology in which genetic and environmental factors are combined with already known ASD risk factors and results of own melatonin research. It leads to the intriguing hypothesis that low melatonin levels, caused by low CYP1A2 activity - which is again early detectable by coffee drinking behaviour-are a risk factor for ASD as well as ID in their offspring. If so, it is more intriguing that this may lead to policies to detect future parents who are at risk and to treatment strategies to reduce ASD and ID risk.

## Acknowledgments

This study was supported by [blinded for review] for contributions in laboratory measurements.

## Financial Disclosures

All authors report no biomedical financial interests or potential conflicts of interest.

All procedures performed in studies involving human participants were in accordance with the ethical standards of the institutional and/or national research committee and with the 1964 Helsinki declaration and its later amendments or comparable ethical standards.

Informed consent was obtained from all individual participants included in the study.

